# Quantifying and Visualizing the Invisible Hazard of Infant Rebreathing

**DOI:** 10.64898/2026.11.06.764821

**Authors:** David Greenblatt, Nadav Itzhak

## Abstract

**BACKGROUND:** Rebreathing of expired gases during prone infant sleeping contributes to hypoxic and hypercapnic exposure, even without airway obstruction. We investigated how anatomical, physiological, and environmental factors influence aerodynamic rebreathing.

**METHODS:** A dynamically scaled in vitro model simulated prone nasal breathing corresponding to infants weighing 2–5 kg, with nostril diameters of 3.2–6.0 mm and breathing frequencies of 25–40 breaths/min. Ambient temperature and sleep-surface characteristics were varied. Rebreathing was quantified using tracer measurements, while flow visualization revealed the underlying aerodynamic mechanisms.

**RESULTS:** Rebreathing exceeded 60% despite unobstructed nasal openings. Increasing ambient temperature from 23 to 30°C increased rebreathing by 68–77%, while reducing simulated infant mass from 5 to 2 kg increased rebreathing by 39–46%. Fibrous surfaces increased rebreathing, particularly near inclined nasal openings. A novel finding was that increasing nostril diameter from 3.2 to 6.0 mm increased rebreathing by up to 50%. Flow visualization revealed vortical structures that retained expired fluid near the nasal openings, facilitating re-inspiration.

**CONCLUSIONS:** Aerodynamic rebreathing depends strongly on simulated infant characteristics and sleeping environments. These findings are consistent with established safe-sleep recommendations and identify nostril diameter, for the first time, as a potentially important anatomical factor warranting clinical investigation.

**IMPACT:**

- Substantial rebreathing can occur during simulated prone infant respiration despite unobstructed nasal openings, revealing an otherwise invisible aerodynamic mechanism of respiratory exposure.
- Variations in simulated infant mass, ambient temperature, and sleep-surface characteristics produce changes in rebreathing consistent with established SIDS risk factors.
- A novel finding of this study is that nostril diameter strongly influences aerodynamic rebreathing, with increases of up to 50% over the tested diameter range.
- These findings provide a physical basis for understanding expired-gas retention and motivate clinical studies investigating the role of nostril geometry in rebreathing susceptibility.

## 1 Introduction

Sudden infant death syndrome (SIDS) remains an important cause of postneonatal mortality despite substantial reductions following supine sleep recommendations [1]. According to the triple-risk model, SIDS may occur when an infant with an intrinsic physiological vulnerability encounters an external stressor during a critical developmental period [1]. Established environmental risk factors include prone sleeping, soft bedding, surface sharing, and overheating [1,2]. A recent analysis of 7,595 sudden unexpected infant deaths (SUID) found that at least 76% involved multiple unsafe-sleep factors [3].

One such respiratory stressor is the rebreathing of expired air containing elevated CO_2_ and reduced O_2_ concentrations. In vulnerable infants, abnormalities in medullary serotonergic signaling may impair protective responses, such as compensatory ventilation or arousal from sleep [4,5]. Previous studies demonstrated substantial CO_2_ accumulation during prone breathing, particularly on soft or porous bedding [6–8], and confirmed increased inspired CO_2_ accompanied by O_2_ depletion in sleeping infants [9,10]. However, close contact with bedding often makes it difficult to distinguish rebreathing from gas retention within porous materials or direct airway obstruction. Rebreathing may therefore represent an external stressor to which vulnerable infants cannot respond adequately.

In our previous study [11], we introduced an aerodynamic approach to infant rebreathing and demonstrated substantial re-inspiration of expired fluid without airway obstruction using a dynamically scaled in-vitro model. By independently varying Reynolds, Strouhal, and densimetric Froude numbers, we isolated the effects of inertia, breathing unsteadiness, and buoyancy. Expired nasal jets impinge on the underlying surface, generating vortical structures that can retain expired fluid near the airways. The present study differs fundamentally in its objectives and scope. Rather than independently varying dimensionless parameters to isolate aerodynamic mechanisms, we systematically vary the underlying anatomical, physiological, and environmental parameters, whose combined effects determine rebreathing under more representative infant conditions. Specifically, we investigate the effects of infant mass and associated tidal volume, nostril diameter, breathing frequency, ambient temperature, and sleep-surface characteristics. The resulting simultaneous changes in Reynolds, Strouhal, and densimetric Froude numbers are interpreted within the infant-relevant ranges established previously [11].

The objectives of the present research are to quantify the effects of these parameters on aerodynamic rebreathing, identify the flow mechanisms responsible for expired-gas retention and subsequent re-inspiration, and assess their potential relevance to established SIDS risk factors. Since accumulated expired gases are invisible and difficult to establish retrospectively [1,12], we employ a dynamically scaled colored-water model that combines quantitative rebreathing measurements with direct flow visualization. This approach enables systematic investigation of rebreathing without direct airway obstruction and identifies previously unexplored anatomical factors that may warrant clinical investigation.

## 2 Methods

### 2.1 Experimental model and dynamic similarity

A dynamically scaled in-vitro model described in [11] was used to investigate aerodynamic rebreathing during prone nasal respiration while excluding direct airway obstruction. Experiments were conducted in a 30 × 40 × 30 cm water-filled Plexiglas tank, where three interchangeable nasal modules were connected to independently driven syringe pumps (Figure 1). Two cross-sectional views of a nasal model are shown in Figure 2 (top row), illustrating two external orifices for expiration (representing nostrils) and an internal orifice for inspiration. Separate internal passages prevent direct cross-contamination between expiration and inspiration flows, ensuring that the extracted fluid represents rebreathing from the surrounding environment rather than residual expired fluid within the nasal passages or connecting tubing. Three separate interchangeable nasal modules were printed using Stereolithography (SLA), with average expiration orifice diameters of 3.2, 4.8, and 6.0 mm, and all with a single central 4.5-mm diameter inspiration orifice (Figure 2, bottom row). The quantitative rebreathing measurement procedure, including tracer calibration and correction for the initial fluid volume in the extraction tubing, was identical to that described and employed in our previous study [11]. The required separate expiration and inspiration orifices are justified because expiration produces two viscous jets that interact with the underlying surface, whereas inspiration generates an approximately potential sink flow. Thus, a single central extraction orifice provides a reasonable approximation of the combined suction through both nostrils, although the detailed near-orifice flow is not reproduced.

**Figure 1.**
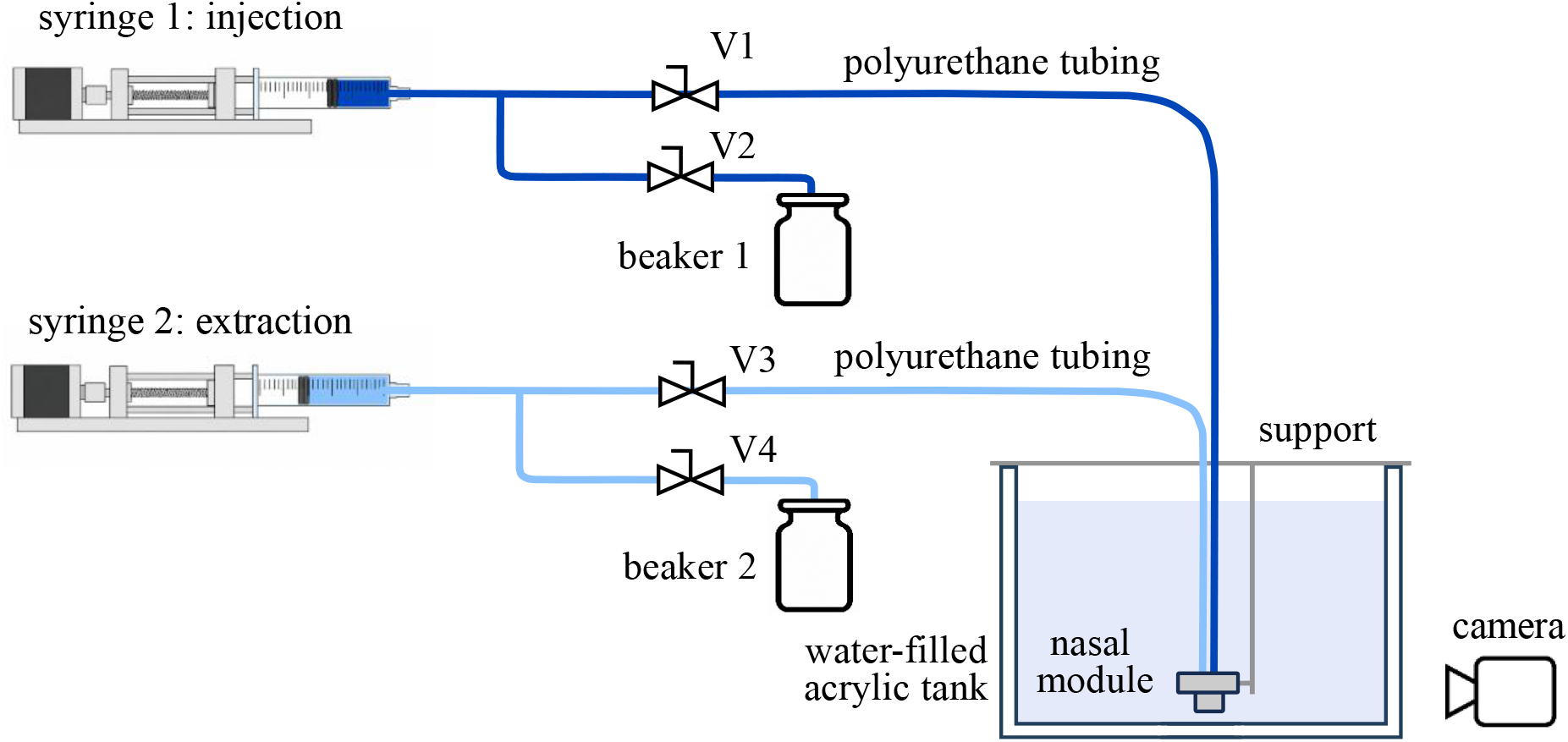
In-vitro water-based experimental setup designed to produce dynamically-scaled Reynolds, Strouhal, and densimetric Froude numbers associated with infant breathing. A detailed description is provided by Itzhak and Greenblatt [11].

**Figure 2.**
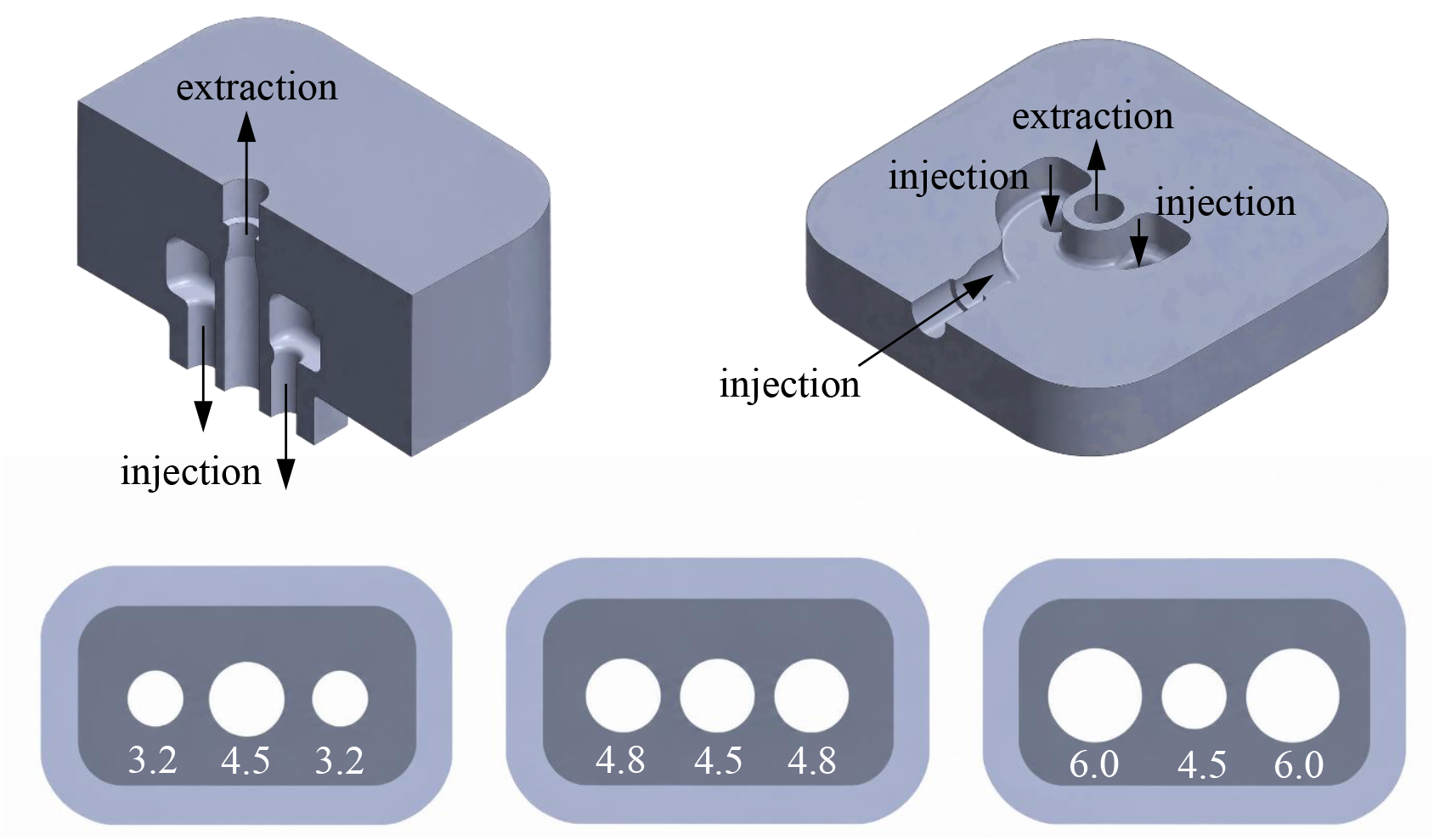
Schematic representations of the nasal modules. Top row: Vertical (left) and horizontal cross-sections of a nasal module. Bottom row: Outer expiration (injection) and central inspiration (extraction) orifice diameters (indicated in mm) for three nasal modules.

Dynamic similarity was based on Reynolds, Strouhal, and inverse squared densimetric Froude numbers, representing the relative contributions of inertia, breathing unsteadiness, and buoyancy, respectively:

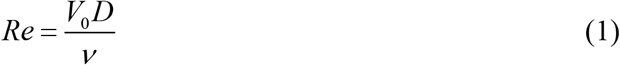

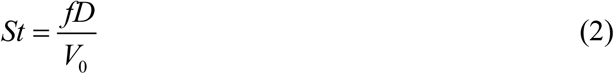

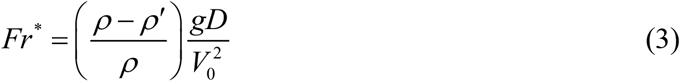

where *V*_0_ is the characteristic expiratory velocity, *D* is the expiration-orifice diameter, *f* is the breathing frequency, *v* is the kinematic viscosity, and *ρ* and *ρ*^*′*^ are the densities of the ambient fluid and expired-fluid analogue, respectively. The signed inverse square of the traditional Froude number (*Fr*^*^ ≡ *Fr*^−2^) is used because it avoids a singularity at neutral buoyancy and distinguishes between positively and negatively buoyant expired fluid. Estimated uncertainties in these dimensionless parameters never exceeded 6%.

### 2.2 Breathing simulation

A periodic breathing waveform was generated by alternately driving the expiration and inspiration syringes. Each breathing period, *T* , was divided into six equal intervals. Expiration accelerated linearly from zero to the prescribed maximum velocity *V*_0_ , remained constant, and then decelerated linearly to zero during the first half-cycle. Inspiration followed the corresponding negative-velocity waveform during the second half-cycle as described in [11]. Infant weight was represented by tidal volume, assuming 7 mL kg^−1^ body mass, consistent with measurements in healthy infants during natural sleep [13]. The principal experiments represented nominal infant masses of 2 and 5 kg, with breathing frequencies of 25 and 40 breaths per minute (bpm), selected to encompass relatively slow and typical breathing conditions in infants [13,14].

### 2.3 Rebreathing measurements & Test Conditions

The injected expired-fluid analogue contained water, HCl, ethanol or sugar, and food coloring. Ethanol or sugar was used to control density, HCl served as a passive tracer for quantitative rebreathing measurements, and food coloring was used for flow visualization. Measurements were performed at density differences representing ambient temperatures of 23, 30, and 36°C, assuming 60% relative humidity, with expired air at 36.5°C, saturated with water vapor, and containing between 4% and 8% CO_2_. Two surface conditions were investigated: a smooth, hard, nonporous and impermeable surface (polyvinyl chloride) and a 3-mm-thick fibrous polyester surface. Polyester was selected because its hydrophobic properties allowed it to retain its structure when submerged and because microscopic examination showed a fiber morphology similar to soft woolen bedding (Figure 3, right).

**Figure 3.**
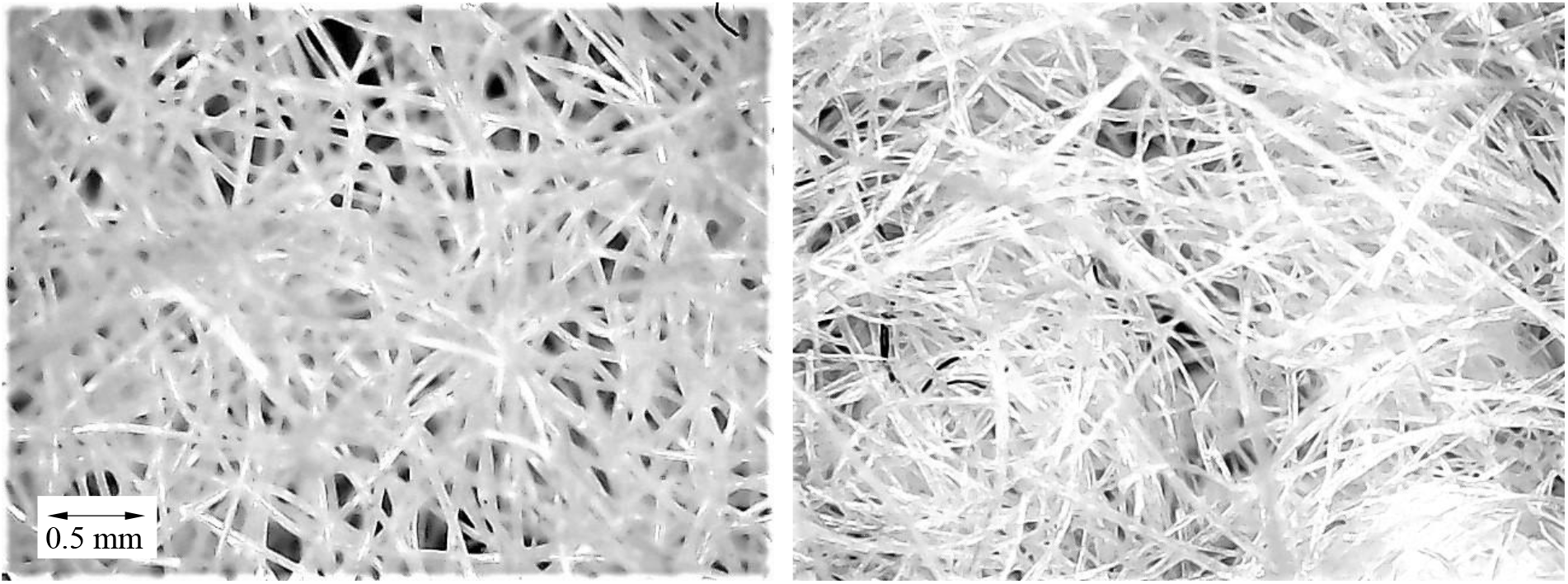
Polyester mat (left) and soft woolen blanket both under 120× magnification.

Before each experiment, the tank and extraction tubing were filled with ambient water and the injection syringe was filled with the calibrated expired-fluid analogue. Alternating expiration and inspiration cycles were performed until the injection syringe was depleted, corresponding to approximately 5–30 breathing cycles depending on the experimental condition. The fluid extracted through the inspiration passage was then collected and its electrical conductivity measured.

The measured conductivity was corrected for ambient water initially contained within the extraction tubing according to

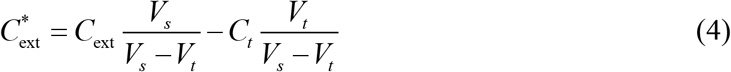

where *C*_ext_ is the measured conductivity of the extracted fluid, *C*_*t*_ is the conductivity of the ambient tank water, and *V*_*s*_ and *V*_*t*_ are the syringe and tubing volumes, respectively. Each set of 5–30 breathing cycles was repeated up to eight times to ensure that each reported average represented at least 40 breathing cycles. Finally, the corrected conductivity was converted to HCl concentration using the calibration relationship and the rebreathing ratio was calculated according to

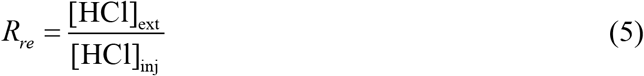

where [HCl]_ext_ and [HCl]_inj_ are the tracer concentrations in the extracted and injected fluids, respectively. The experimental matrix included systematic variation of ambient-temperature analogue, nominal infant mass/tidal volume, nostril diameter, breathing frequency, and surface type. The principal conditions are summarized in Table 1.

**Table 1.** Summary of the experimental conditions considered in this study.

| **Parameter** | **Conditions** |
|---|---|
| expiration-orifice diameter, *D* | 3.2, 4.8, 6.0 mm |
| inspiration-orifice diameter | 4.5 mm |
| nominal infant mass | 2 and 5 kg |
| tidal volume | 7 mL kg<sup>−1</sup> |
| breathing frequency | 25 and 40 bpm |
| ambient-temperature analogue | 23, 30, 36°C |
| assumed expired CO<sub>2</sub> concentration | 4% to 8% |
| assumed ambient relative humidity | 60% |
| surface type | Smooth, hard, nonporous; 3-mm fibrous polyester |
| Reynolds number, *Re* | 121 to 373 |
| Strouhal number, *St* | 1.17×10<sup>−3</sup> to 7.71×10<sup>−3</sup> |
| Froude number, *Fr*<sup>\*</sup> | –6.30×10<sup>−3</sup> to +1.74×10<sup>−2</sup> |
| quantitative repetitions | 3 to 8 per condition |
| breathing cycles per measurement | 5 to 30 |
| primary outcome | Rebreathing ratio, *R<sub>re</sub>* (%) |

## 3 Results and Discussion

### 3.1 Ambient Temperature Effects

The effects of both ambient temperature and infant body mass (tidal volume) on rebreathing are presented in Figure 4, with the corresponding Reynolds, Strouhal and Froude numbers indicated in Table 2. Note that changes in body mass affect all three dimensionless parameters, whereas changes in ambient temperature primarily affect the Froude number.

**Table 2.** Dimensionless parameters corresponding to different cases presented in Figure 4.

| No. | *m* (kg) | *T*<sub>amb</sub> (°C) | *Re* | *St* (10<sup>−3</sup>) | *Fr*<sup>\*</sup> (10<sup>−3</sup>) |
|---|---|---|---|---|---|
| 1 | 5 | 23 | 362 | 1.6 | +1.3 |
| 2 | 2 | 23 | 145 | 4.0 | +8.4 |
| 3 | 5 | 30 | 362 | 1.6 | +0.58 |
| 4 | 2 | 30 | 145 | 4.0 | +3.6 |
| 5 | 2 | 36 | 152 | 4.0 | −0.60 |

**Figure 4:**
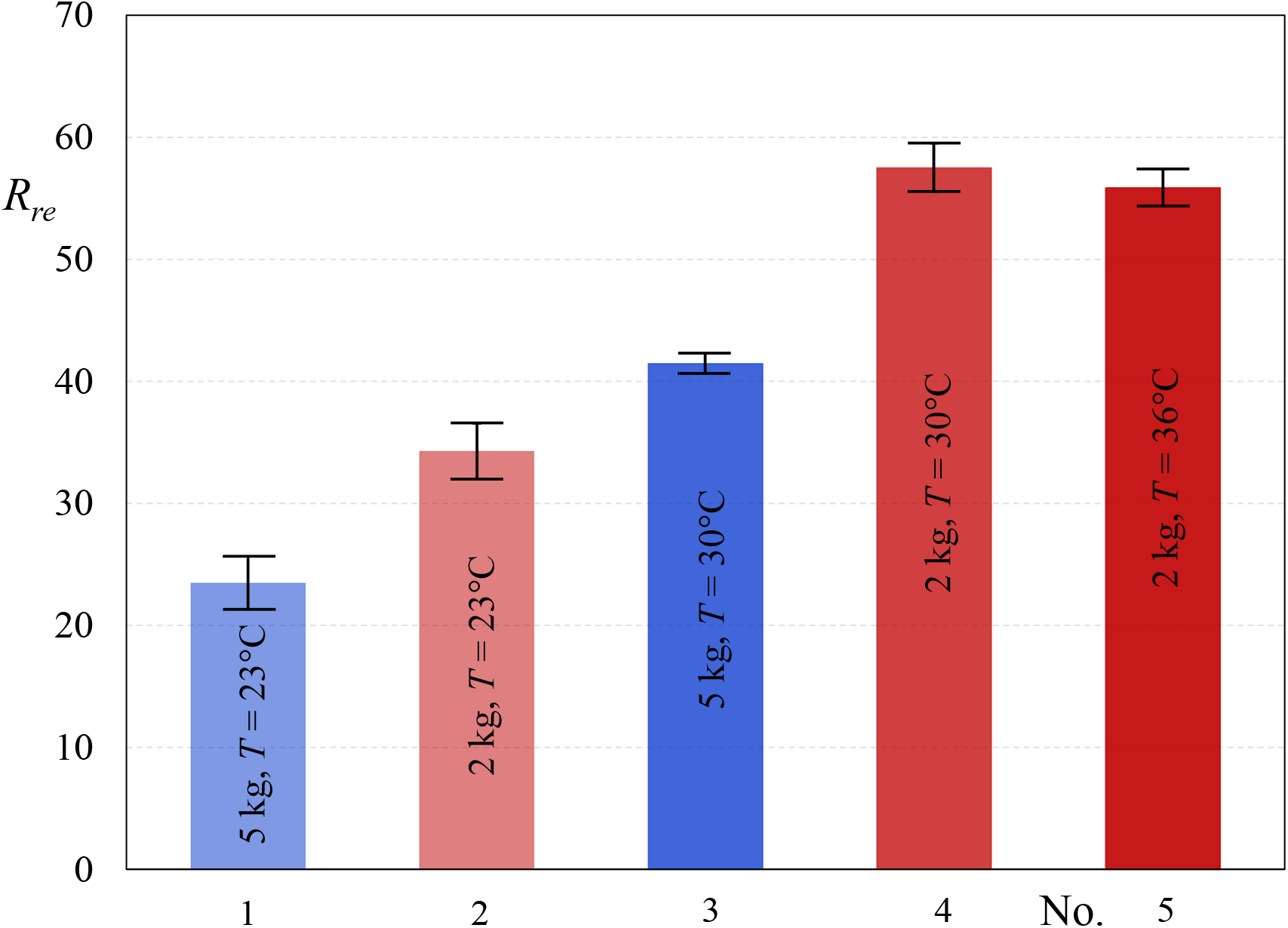
Percentage rebreathing for infants weighing 2 and 5 kg, breathing at 25 bpm, at ambient temperatures of 23, 30 and 36°C, with standard deviations indicated (*D* = 4.8 mm, *H* = 6.4 mm, *H* / *D* = 1.33). Dimensionless groups are shown in Table 2.

The effect of the ambient temperature is consistent, irrespective of the infant size, where an increase from 23 to 30°C substantially increases rebreathing by 77% and 68% for simulated infant masses of 5 and 2 kg, respectively. A further increase in ambient temperature from 30 to 36°C maintains the elevated rebreathing ratio within experimental uncertainty. Expired fluid corresponding to 23°C (large positive Froude numbers) rises away from the orifices due to buoyancy, thereby reducing the fraction of expired fluid that is rebreathed during subsequent inspiration (see discussion below). Epidemiological studies have reported an association between elevated ambient temperatures and increased SIDS risk, particularly during summer months [15,16]. The present experimental results are consistent with these observations, demonstrating that elevated ambient temperatures can increase rebreathing by modifying the buoyancy of expired gases and promoting their accumulation near the nostrils. Although this agreement does not establish a causal relationship between rebreathing and SIDS, it provides a plausible aerodynamic mechanism that is consistent with the observed epidemiological trends.

The effect of simulated infant mass from 5 to 2 kg is also consistent, irrespective of the ambient temperature, with rebreathing increases of 46% and 39%, for ambient temperatures of 23 to 30°C, respectively. These observations are explained by the simultaneous decrease in Reynolds number (*Re* = 362 to 145) and increase in Strouhal number *St* = 1.6×10^−3^ to 4.0×10^−3^) , both of which were previously shown to increase rebreathing [11]. The accompanying increase in positive buoyancy (*Fr*^*^), which tends to reduce rebreathing by transporting expired fluid away from the orifices, opposes these effects. The present results therefore suggest that the combined effects of Reynolds and Strouhal numbers dominate the opposing influence of buoyancy. As in the case of ambient temperature effects discussed above, epidemiological studies have consistently identified low birth weight, restricted fetal growth, and poor postnatal weight gain as risk factors for SIDS [17–20]. These observations are consistent with the present experiments, which demonstrate substantially greater rebreathing for a simulated infant mass of 2 kg compared with 5 kg. Again, our experiments do not establish a causal relationship between infant mass, rebreathing, and SIDS. But they do identify an aerodynamic mechanism that may contribute to the observed epidemiological associations.

The flow visualizations in Figure 5 provide insight into the mechanisms underlying the rebreathing measurements in Figure 4. (Note that for these visualizations, only the first breathing cycle is shown in order to avoid excessive contamination produced by the coloring.) For the 5-kg infant at 23°C (*R*_*re*_ = 23.5%), the expired fluid impinges on the surface and forms rolled-up vortices that spread laterally during expiration (Figure 5a,b). At the middle of the breathing cycle (Figure 5c, top row), positive buoyancy causes the expired fluid to rise around the sides of the module and away from the inspiration orifice, thereby limiting subsequent rebreathing. In contrast, for the 2-kg infant at 30°C (*R*_*re*_ = 57.5%), the expired fluid forms more compact vortical structures that remain concentrated beneath the module. At the corresponding instant (Figure 5c, bottom row), considerably less fluid has risen away from the module, leaving a substantial accumulation near the inspiration orifice that is subsequently drawn in during inspiration (Figure 5d). These observations are consistent with the decrease in Reynolds number (*Re* = 362 to 145), which promotes the accumulation of expired fluid closer to the orifices, and the increase in Strouhal number (*St* = 1.6×10^−3^ to 4×10^−3^), which reduces the time available for the vortical structures to move away before inspiration [11]. Although positive buoyancy is approximately 6 times larger for the smaller simulated infant, its influence is insufficient to overcome the combined effects of Reynolds and Strouhal numbers, resulting in substantially greater rebreathing. Visualization of rebreathing is shown dramatically in Supplementary Video 1, which provides a direct visualization of the otherwise invisible aerodynamic mechanisms of rebreathing.^*^ The additional increase in ambient temperature from 30 to 36°C maintains the elevated rebreathing ratio because the very slightly negatively buoyant expired fluid tends to remain near the orifices (not shown). Thus, both high temperature conditions produce substantial rebreathing, but via different buoyancy mechanisms.

**Figure 5:**
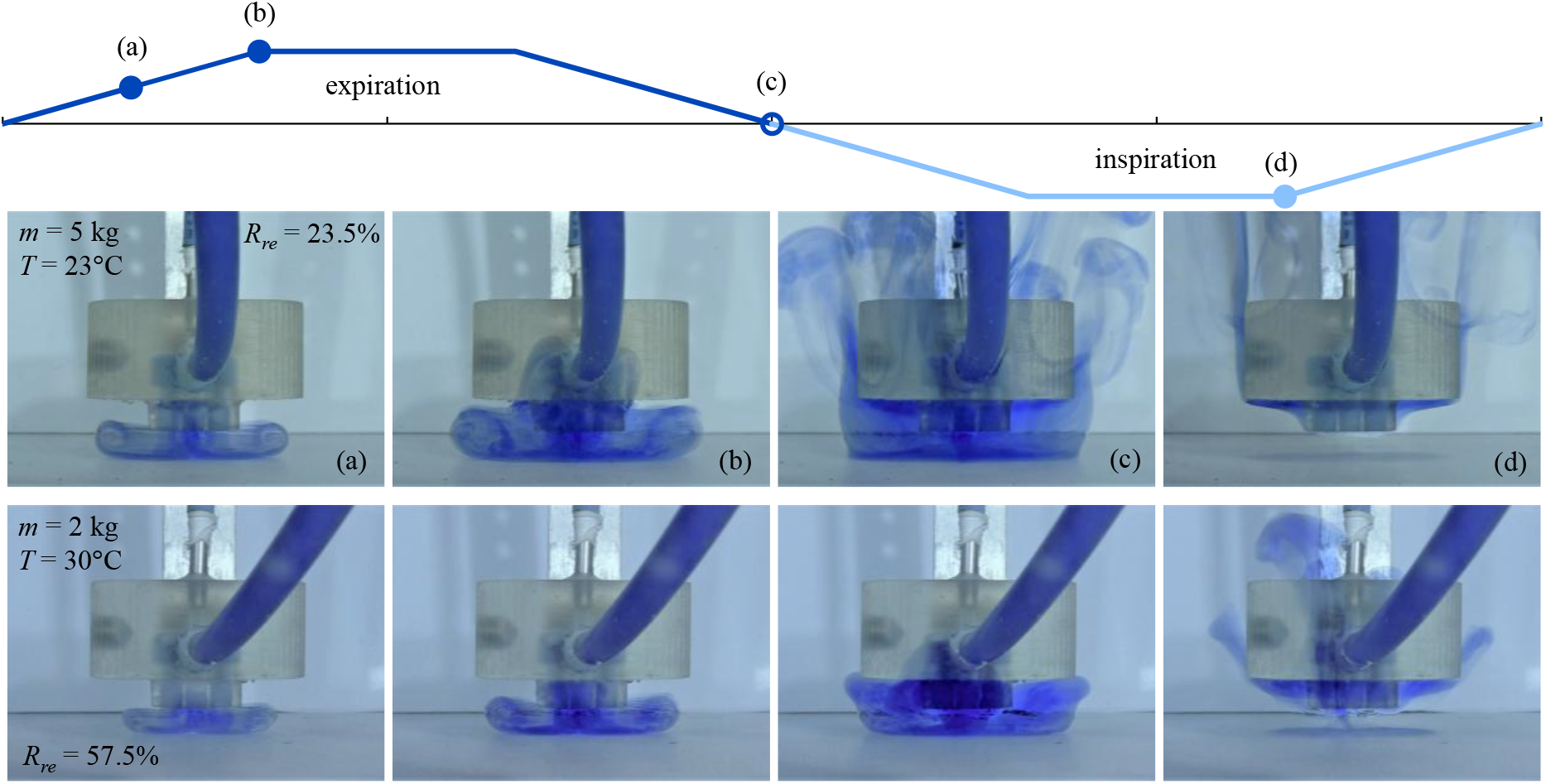
Front view flow visualization for two examined cases from Figure 4. Top: simulated breathing cycle where (a)-(d) indicate the phases of the images. Top row: simulated infant mass of 5 kg and ambient temperature of 23°C. Bottom row: simulated infant mass of 2 kg and ambient temperature of 30°C.

### 3.2 Sleep-Surface Characteristics

The effects of sleep-surface characteristics on aerodynamic rebreathing were investigated by comparing a smooth, impermeable surface with a 3-mm-thick polyester fiber mat. The latter was selected because its fibrous structure resembles that of a soft woolen bedding (Figure 3), which has previously been associated with increased SIDS risk [1,21,22]. Unlike conventional bedding studies, the nasal openings remained unobstructed, thereby isolating aerodynamic rebreathing from direct airway obstruction. Experiments were performed for the conditions *m* = 2 kg (*∀* = 14 mL), *D* = 3.2 mm, and 25 bpm (*Re* = 228 , *St* = 1.2×10^−3^) , with Froude numbers close to zero (| *Fr*^*^ |< 10^−4^) representing neutrally buoyant expired air.

Figure 6 compares the flow development over the polyester fiber mat and the smooth, impermeable surface, with the expiration orifices positioned 6.4 mm above the surface. The rebreathing ratio increases from 44.5% over the smooth surface to 54% over the fibrous mat, corresponding to a relative increase of approximately 21%. Over the smooth surface, the expired fluid impinges on the surface and spreads laterally, forming well-defined counter-rotating vortices. In contrast, the fibrous mat restricts lateral spreading, while some of the expired fluid penetrates its porous structure. Hence, the fibrous layer acts as a temporary reservoir of expired fluid, part of which can subsequently be recovered during inspiration. However, fluid retained within the layer may not be readily extracted, which could explain the relatively modest increase in rebreathing.

**Figure 6:**
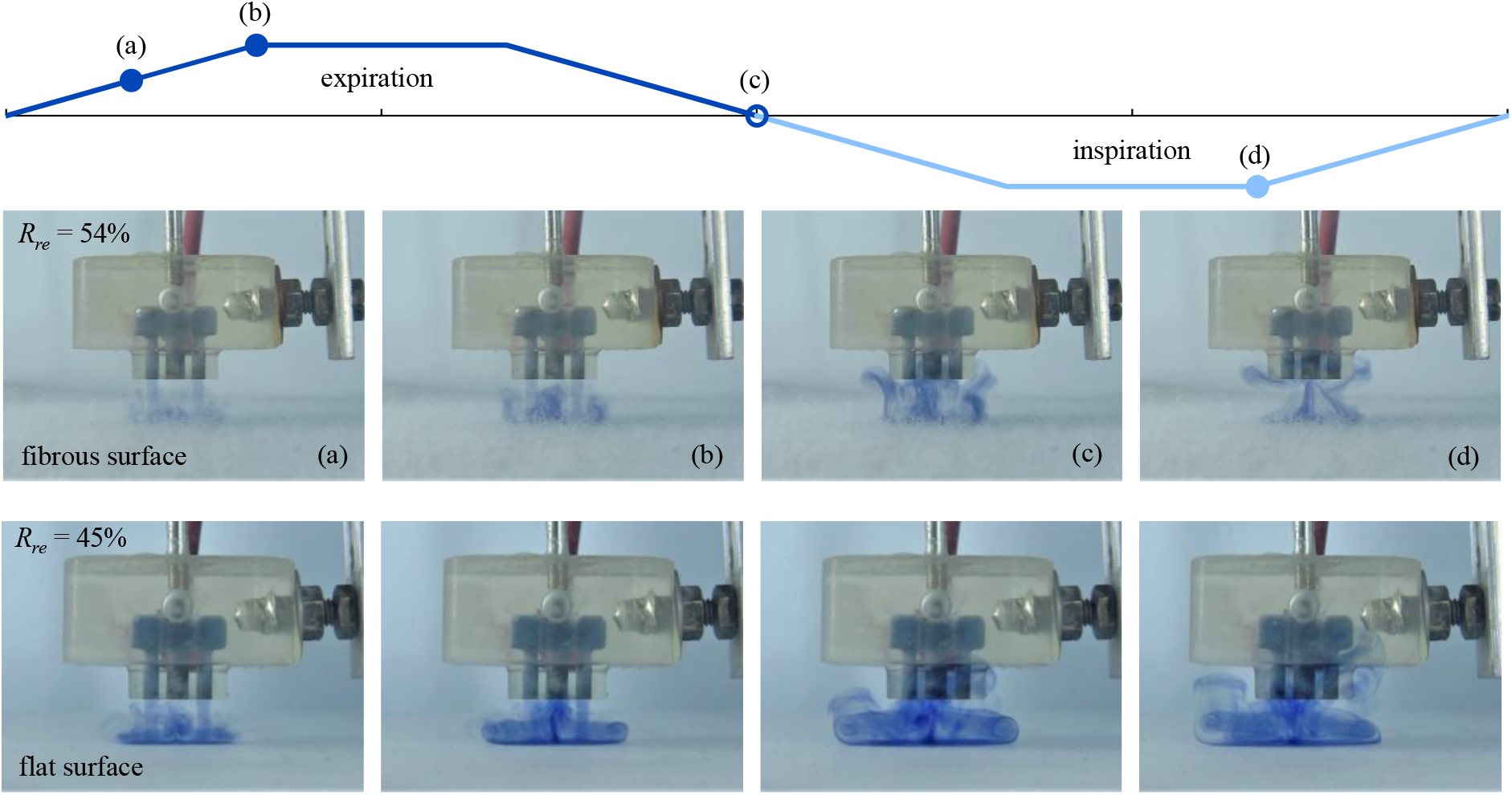
Flow visualization illustrating the effect of the fibrous surface with the nasal module oriented normal to the surface. Experimental conditions: *m* = 2 kg (*∀* = 14 mL), *D* = 3.2 mm, and 25 bpm.

The relevance of these observations to de facto soft woolen bedding lies primarily in the convective transport mechanisms. In both cases, the fibrous structure modifies the impinging expiratory flow, while some expired fluid penetrates and remains temporarily within the interconnected pore spaces. However, molecular diffusion of CO_2_ in air is substantially faster than dye diffusion in water, even if the permeability, porosity, and fluid–fiber interactions are similar. Consequently, although convection dominates expired-fluid transport above the surfaces, the relative importance of convection and diffusion within the fibrous layers may differ considerably. The present observations therefore provide qualitative evidence of a possible retention-and-exchange mechanism, but do not establish quantitative equivalence between the polyester mat in water and woolen bedding in air. Direct measurements of CO_2_ rebreathing over representative bedding materials would be required to assess the magnitude of this effect under physiological conditions.

The influence of the fibrous surface becomes substantially more pronounced when the nasal module is inclined at 30° to the horizontal and placed directly above the surface (Figure 7). Under these conditions, the rebreathing ratio increases from 11% over the smooth surface to 34% over the fibrous mat, representing an approximately threefold increase. As observed previously, the fibrous layer restricts lateral transport and allows some expired fluid to penetrate its porous structure. However, the inclined flat configuration introduces a preferential direction for the expiratory flow, resulting in pronounced lateral convection over the smooth surface and transporting much of the expired fluid away from the nasal module. This is clearly evident in both the side- and front-view visualizations. In contrast, the fibrous mat inhibits this convective removal, leaving considerably more expired fluid near the inspiration orifice. Furthermore, the reduced clearance between the module and the surface increases the interaction between the expiratory flow and the bedding, potentially enhancing expired-fluid retention. The combined effects of inclination and reduced clearance therefore accentuate the differences between the two surfaces. Although the relative contributions of these geometric changes cannot be determined independently, the results demonstrate that bedding characteristics can have a substantially greater influence on aerodynamic rebreathing when the nasal module is inclined and positioned close to the surface.

**Figure 7:**
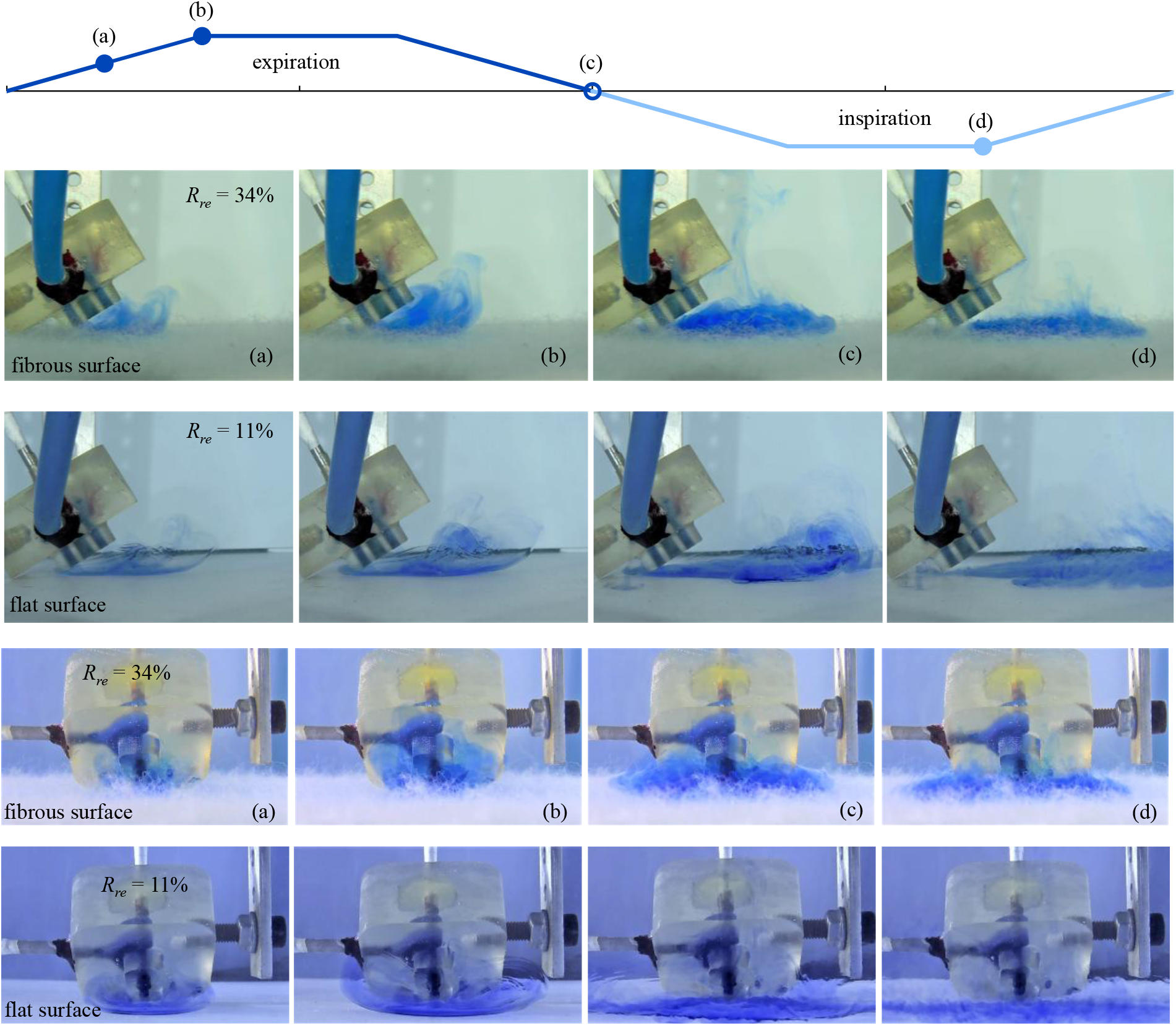
Flow visualization illustrating the effect of a fibrous surface with the nasal module oriented at 30° to the surface. Top two rows: front view; bottom two rows: side view. Experimental conditions: *m* = 2 kg (*∀* = 14 mL), *D* = 3.2 mm, and 25 bpm.

### 3.3 Nostril Diameter and Breathing Frequency

The effects of nostril diameter and breathing frequency on rebreathing are presented in Figure 8. Increasing the mean nostril diameter from 3.2 to 6 mm increased the rebreathing ratio by up to 50%, reaching approximately 65%. Conversely, increasing breathing frequency from 25 to 40 bpm reduced rebreathing by up to approximately seven percentage points. These trends were observed for both buoyancy conditions investigated. Increasing nostril diameter simultaneously decreased Reynolds number and increased Strouhal number (Table 3), favoring retention of expired fluid near the nostrils. In contrast, increasing breathing frequency at constant tidal volume increased the Reynolds number while leaving Strouhal number approximately unchanged, thereby promoting convective transport away from the nostrils.

**Table 3:** Corresponding dimensionless parameters for all cases presented in Figure 8, for *m* = 2 kg (*∀* = 14 mL).

| *D* (mm) | *f* (bpm) | *T* (°C) | CO<sub>2</sub> (%) | *Re* | *St* (10<sup>−2</sup>) | *Fr*<sup>\*</sup> (10<sup>−2</sup>) |
|---|---|---|---|---|---|---|
| 3.2 | 25 | 27 | 4 | 227 | 0.117 | +0.075 |
| 3.2 | 40 | 27 | 4 | 363 | 0.117 | +0.029 |
| 4.8 | 25 | 27 | 4 | 151 | 0.395 | +0.57 |
| 4.8 | 40 | 27 | 4 | 242 | 0.395 | +0.22 |
| 6.0 | 25 | 27 | 4 | 121 | 0.771 | +1.74 |
| 6.0 | 40 | 27 | 4 | 193 | 0.771 | +0.68 |
| 3.2 | 25 | 34 | 8 | 233 | 0.117 | −0.027 |
| 3.2 | 40 | 34 | 8 | 373 | 0.117 | −0.011 |
| 4.8 | 25 | 34 | 8 | 156 | 0.395 | −0.21 |
| 4.8 | 40 | 34 | 8 | 249 | 0.395 | −0.081 |
| 6.0 | 25 | 34 | 8 | 124 | 0.771 | −0.63 |
| 6.0 | 40 | 34 | 8 | 199 | 0.771 | −0.25 |

**Figure 8:**
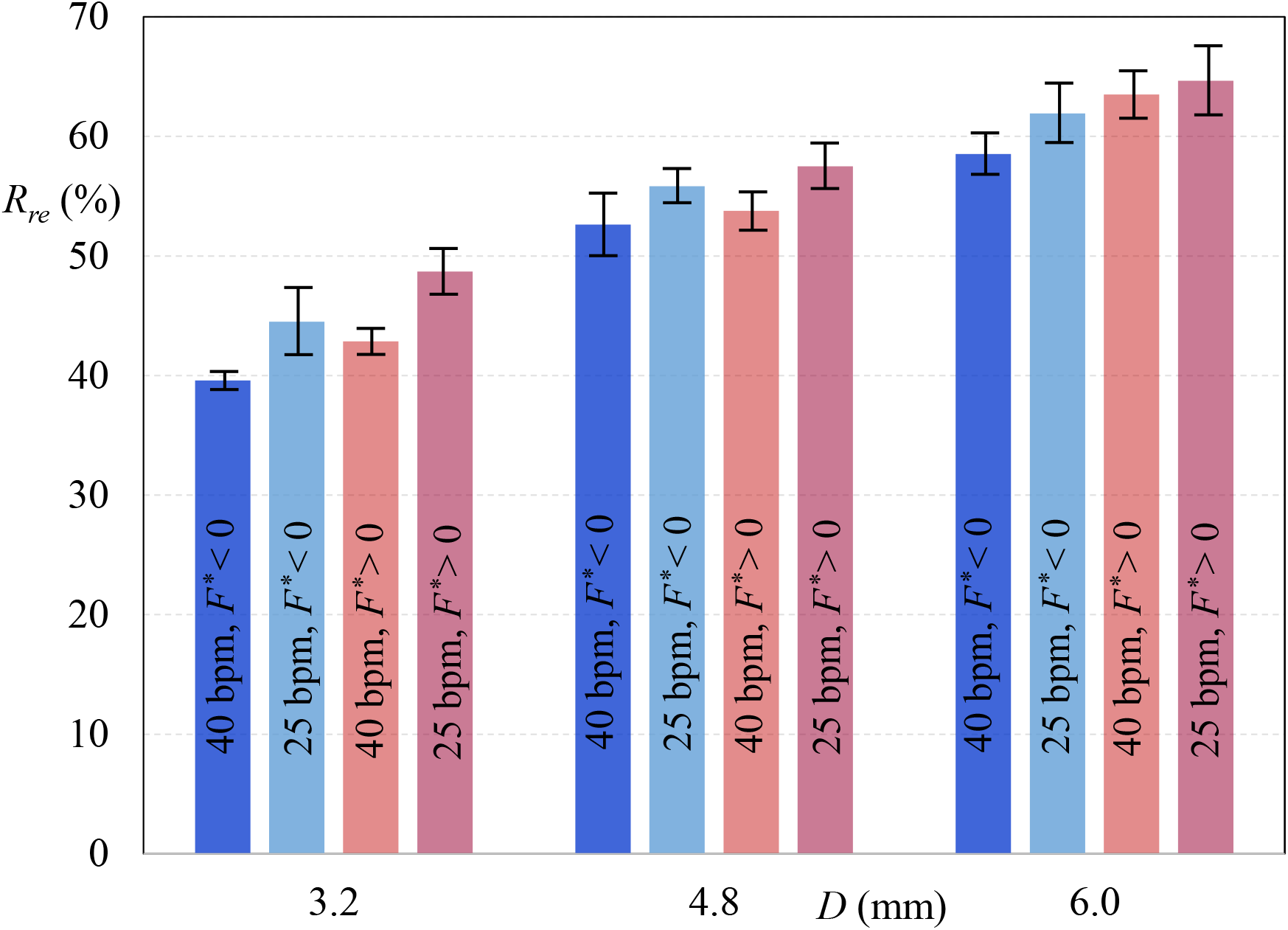
Rebreathing percentages as a function of nostril diameter for *m* = 2 kg (*∀* = 14 mL) and breathing frequencies of 25 and 40 bpm for different expired air CO_2_ content (see Table 3), with standard deviations indicated.

Front and side view flow visualizations (Figure 9) provide further insight into the effects of nostril diameter. The smaller orifices generated more extensive secondary vortical structures that transported expired fluid laterally and vertically away from the surface. These structures appeared to deform and break up, promoting dispersion and mixing of the expired fluid before subsequent inspiration. This behavior is consistent with Reynolds-number effects on secondary vortex formation and transport. Furthermore, the lower Strouhal number associated with smaller nostrils provides more time for the vortical structures to convect away before subsequent inspiration. Thus, the combined effects of Reynolds number on secondary vortex formation and Strouhal number on vortex transport provide a plausible explanation for the observed reduction in rebreathing with decreasing nostril diameter.

**Figure 9:**
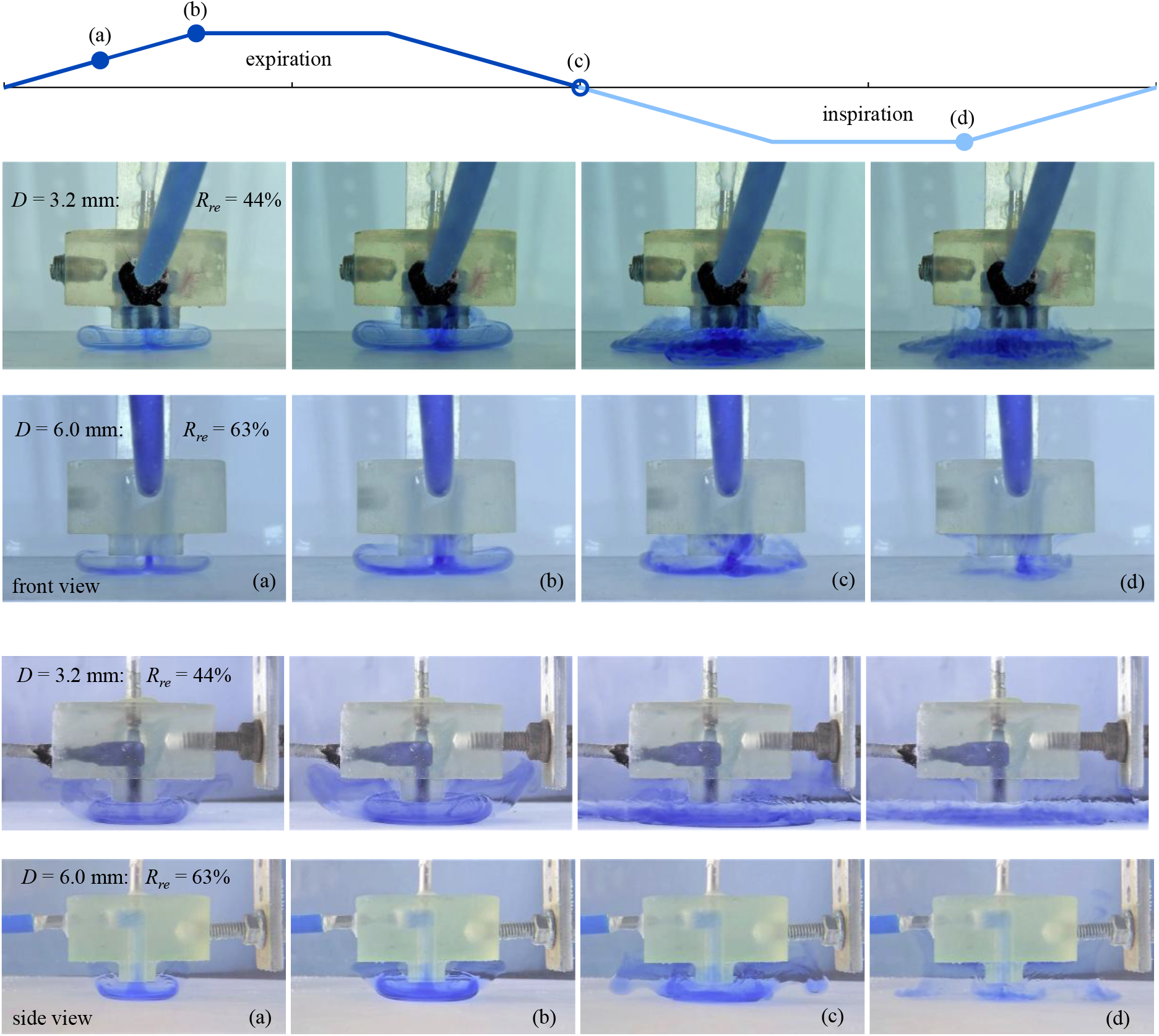
Flow visualization comparison for orifice diameters of 3.2 mm and 6.0 mm, respectively, for *m* = 2 kg (*∀ =*14 mL) and 25 bpm. Top two rows: front view; bottom two rows: side view.

These findings identify nostril diameter as a potentially important, but previously unexplored, anatomical parameter governing aerodynamic rebreathing. This raises the possibility that anatomical variability may contribute to differences in respiratory exposure among infants. For example, anthropometric studies have identified differences in external nasal dimensions between population groups, including greater mean nose breadth in Black than White newborn infants [23]. Marked differences in SIDS and SUID incidence have also been reported between other racial and ethnic groups [24]. However, external dimensions do not necessarily reflect effective nostril diameter, and no direct relationship between infant nostril dimensions and SIDS incidence has been established. The present results suggest that such a relationship warrants investigation, particularly through measurements of individual nostril geometry and corresponding aerodynamic rebreathing. Any potential anatomical contribution would need to be evaluated alongside established physiological, environmental, and socioeconomic risk factors.

The present findings suggest two complementary approaches for assessing the potential clinical significance of nostril geometry in infant rebreathing. First, noninvasive postnatal screening using calibrated photographs could provide measurements of nostril area, wetted perimeter, and hence the hydraulic diameter. Combined with infant mass and breathing frequency, these measurements could be used to estimate the governing aerodynamic parameters and, using an experimentally calibrated relationship, predict susceptibility to rebreathing under prone sleeping conditions. Second, retrospective post-mortem measurements could determine whether nostril geometry and predicted rebreathing susceptibility differ systematically between SIDS cases and appropriately matched controls, accounting for established risk factors and possible post-mortem anatomical changes. Previous studies have identified associations between craniofacial morphology and SIDS or sudden unexpected infant death [25,26], although the specific role of nostril hydraulic diameter remains unknown. Retrospective findings could motivate prospective studies to assess the predictive value of these anatomical measurements.

## 4 Physiological Relevance and Clinical Implications

The present results demonstrate that substantial rebreathing can occur during simulated prone respiration despite unobstructed nasal openings, with rebreathing ratios exceeding 60% under certain conditions. Measurements in sleeping infants have previously demonstrated that rebreathing expired gases from bedding can simultaneously increase inspired CO_2_ and reduce inspired O_2_ concentrations [9]. Although gas concentrations were not measured directly in the present experiments, the potential magnitude of inspired CO_2_ exposure can be estimated using a simple mixing model:

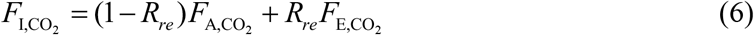

Where 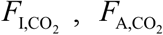 , and 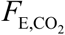 are the inspired, ambient, and expired CO_2_ fractions, respectively. Assuming ambient and expired CO_2_ concentrations of 0.04% and 4%, respectively, rebreathing ratios of 40% and 60% correspond to estimated inspired CO_2_ concentrations of approximately 1.6% and 2.4%. These estimates assume uniform mixing and constant expired CO_2_ concentration; they illustrate potential exposure rather than measured physiological conditions.

A potentially important consequence of sustained rebreathing is a positive-feedback mechanism. If increased inspired CO_2_ is accompanied by inadequate alveolar ventilation, arterial and subsequently expired CO_2_ concentrations may increase, further elevating inspired CO_2_ during subsequent breaths. For example, at a rebreathing ratio of 60%, increasing expired CO_2_ from 4% to 6% would increase the estimated inspired concentration from approximately 2.4% to 3.6%. Normally, hypercapnia and hypoxia stimulate compensatory ventilation and arousal. However, impaired ventilatory regulation, arousal, or autoresuscitation may compromise these protective responses [4,27]. This feedback mechanism is physiologically plausible but was not demonstrated experimentally, and its progression would depend on metabolic CO_2_ production, effective alveolar ventilation, and physiological compensation.

The interaction between aerodynamic rebreathing and impaired protective responses is consistent with the triple-risk model of SIDS, in which an underlying vulnerability interacts with an external stressor during a critical developmental period [4]. Our findings identify aerodynamic mechanisms that can produce substantial rebreathing without direct airway obstruction and demonstrate their sensitivity to infant characteristics, nostril geometry, ambient temperature, and sleep-surface properties. These findings provide a possible physical explanation for respiratory hazards associated with prone sleeping and unfavorable sleep environments.

The observed accumulation of expired fluid near the nasal openings also suggests that ambient air movement could reduce rebreathing by dispersing CO_2_-rich gases before they are re-inspired. This mechanism is consistent with the epidemiological association between fan use during sleep and reduced SIDS risk reported by Coleman-Phox et al. [28]. Although ventilation was not varied in the present experiments, the flow visualizations provide a plausible aerodynamic explanation for this association.

Although the present experiments do not establish a causal relationship between aerodynamic rebreathing and SIDS, the findings are consistent with established recommendations for supine sleeping on firm, flat surfaces without soft bedding [1]. The newly identified influence of nostril diameter further motivates clinical investigation of whether nasal geometry contributes to individual susceptibility to rebreathing.

## 5 Conclusions

The present study demonstrates that substantial aerodynamic rebreathing can occur during simulated prone infant respiration despite unobstructed nasal openings, with rebreathing ratios exceeding 60% under certain conditions. Unlike our previous investigation, which independently varied the governing dimensionless parameters, the present study systematically examined the combined aerodynamic effects of clinically relevant anatomical, physiological, and environmental parameters.

Rebreathing increased with decreasing simulated infant mass, increasing ambient temperature, larger nostril diameter, and fibrous sleep surfaces. Flow visualization revealed how changes in jet dynamics, vortex formation, and buoyancy influence the accumulation and subsequent re-inspiration of expired fluid. The effects of infant mass (i.e., tidal volume), ambient temperature, and surface characteristics are consistent with established epidemiological risk factors for SIDS, providing a physical explanation for their potential influence on rebreathing.

A particularly important and novel finding is the strong dependence of rebreathing on nostril diameter. Increasing the diameter from 3.2 to 6.0 mm increased rebreathing by up to 50%, identifying nostril geometry as a previously unexplored anatomical factor that may influence susceptibility to rebreathing. This finding warrants retrospective anatomical comparisons between SIDS cases and controls, followed by prospective clinical investigations.

Although these in-vitro experiments do not establish a causal relationship between aerodynamic rebreathing and SIDS, they demonstrate physical mechanisms capable of producing substantial re-inspiration of expired fluid without airway obstruction. The findings are consistent with established safe-sleep recommendations and identify nostril geometry as a potentially important new consideration in understanding infant rebreathing.

## Ethics Approval

This study involved experiments using a dynamically scaled *in vitro* physical model and did not involve human participants, human biological materials, or animals. Institutional ethics approval and informed consent were therefore not required.

## Funding

N.I. acknowledges financial support from the Leonard and Diane Sherman Interdisciplinary Graduate School Fellowship at the Technion–Israel Institute of Technology.

## Competing Interests

The authors declare no competing interests.

## Data Availability

The data supporting the findings of this study are available from the corresponding author upon request.

## Author Contributions

D.G. conceived and designed the study, supervised the research, interpreted the results, and wrote the majority of the manuscript. N.I. conducted the experiments, performed the data analysis, and contributed to the interpretation of the results and manuscript preparation. Both authors reviewed and approved the final manuscript.

## Footnotes

* Supplementary Video 1. Flow visualization of aerodynamic rebreathing under simulated prone infant respiration. Available on YouTube: https://www.youtube.com/watch?v=wdO20sLUfAA.

## Notes

https://www.youtube.com/watch?v=wdO20sLUfAA

### Competing Interest Statement

The authors have declared no competing interest.

